# STRATA: Spatial Regulon Field Theory Reveals Coupling Architecture of Human Skin and Its Homogenization in Melanoma

**DOI:** 10.64898/2026.02.24.707661

**Authors:** Jeng-Wei Tjiu

## Abstract

Spatial transcriptomics captures gene expression in tissue context, yet current analyses reduce continuous regulatory landscapes to discrete cell clusters, discarding the geometry of intercellular regulation. Here we introduce STRATA (Spatial Transcription-factor Regulatory Architecture of Tissue Analysis), a differential-geometric framework that constructs continuous regulon activity fields from transcript coordinates, computes their coupling tensor to quantify local co-regulation between transcription factor programs, and derives a Regulon Stability Index from the Jacobian singular value decomposition. Applied to Xenium in situ data from human skin melanoma (382 genes, 13.7 million transcripts), STRATA identifies coupling phase boundaries — positions where the regulatory logic of tissue changes — that track histological tissue architecture (Pearson r = 0.32 with the dermal–epidermal junction marker KRT-diff, r = 0.51 with maximum principal stretch σ_1_; P < 10^−10^). Within-tissue comparison reveals that the melanoma microenvironment does not abolish regulon coupling but homogenizes it: coupling variance decreases 28% and phase boundary intensity drops 18% relative to the epidermal zone. STRATA transforms spatial transcriptomics from cell cataloguing to continuous field analysis of regulatory tissue architecture.

## Introduction

Spatial transcriptomics technologies now resolve gene expression at subcellular resolution across intact tissue sections. Platforms such as 10x Genomics Xenium, Vizgen MERFISH, and NanoString CosMx detect individual RNA transcripts in situ, producing datasets of millions of molecules with precise spatial coordinates. These data contain rich information about the continuous regulatory landscape of tissue — how transcription factor programs vary, interact, and transition across space. Yet the dominant analytical paradigm reduces this continuous information to discrete outputs: cells are segmented, clustered into types, and mapped back onto tissue coordinates. The regulatory geometry that exists between and across cell boundaries is lost.

This discretization is not merely an aesthetic concern. Tissue function emerges from the spatial organization of regulatory programs — the way transcription factor activities covary locally, how co-regulation structures transition at tissue interfaces, and where regulatory configurations are stable versus labile. These properties are inherently continuous and cannot be captured by cataloguing cell types at discrete positions. What is needed is a framework that treats regulatory programs as continuous spatial fields and provides mathematical tools to characterize their geometry.

Here we introduce STRATA (Spatial Transcription-factor Regulatory Architecture of Tissue Analysis), a three-layer differential-geometric framework for continuous regulon field analysis of spatial transcriptomics data. Layer 1 constructs continuous regulon activity fields from transcript coordinates via kernel density estimation and weighted gene combination. Layer 2 computes a coupling tensor that quantifies the local covariance structure between regulon programs, revealing where and how transcription factor activities co-vary in space. Layer 3 derives a Regulon Stability Index (RSI) from the singular value decomposition (SVD) of the spatial Jacobian, identifying positions where the regulon configuration is rigid versus deformable. Together, these layers define coupling phase boundaries — spatial positions where the regulatory logic of tissue changes — that we show correspond to histologically defined tissue architecture.

We validate STRATA on Xenium in situ data from a human skin melanoma section (382 genes, 13.7 million transcripts) and demonstrate that it reveals a previously unrecognized property of the melanoma microenvironment: the tumor does not abolish regulon coupling but homogenizes it, erasing the structured spatial heterogeneity that characterizes normal tissue architecture.

## Results

### STRATA constructs continuous regulon fields from transcript coordinates

The first layer of STRATA transforms discrete transcript coordinates into continuous regulon activity fields (Fig. 1a). For each gene *g* in the spatial transcriptomics panel, we compute a density field ρ_g(s) at every tissue position s using kernel density estimation (KDE) with a Gaussian kernel of bandwidth h (default 40 μm, approximately two cell diameters). These per-gene density fields are then log-normalized analogously to standard single-cell RNA-seq processing, yielding expression fields x_g(s), and z-scored to produce standardized fields z_g(s) with zero mean and unit variance across the tissue section.

**Figure 1.**
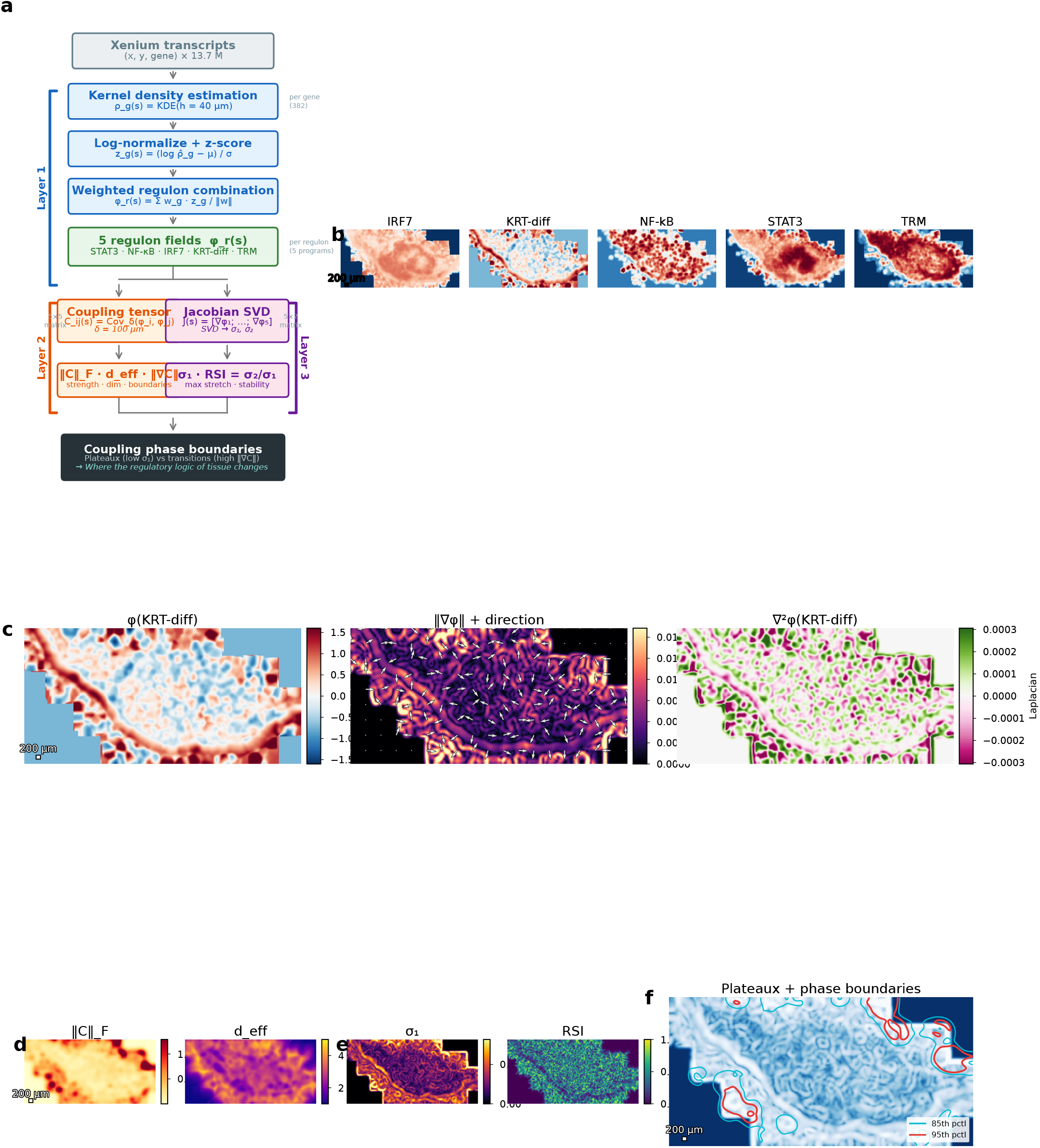
STRATA framework overview and application to human skin melanoma. (a) Pipeline schematic: transcript coordinates are converted to gene density fields via KDE, log-normalized, z-scored, and combined into regulon activity fields. The coupling tensor C(s) captures local co-regulation geometry; the Jacobian SVD yields the stability landscape. (b) Five regulon fields (IRF7, KRT-diff, NF-κB, STAT3, TRM) computed from 13.7 million Xenium transcripts. (c) KRT-diff regulon: activity field (left), gradient magnitude with quiver plot showing gradient direction perpendicular to the epidermal surface (center), and Laplacian (right). (d) Coupling strength ⍰C⍰_F (left) and effective dimensionality d_eff (right). (e) Maximum principal stretch σ_1_ (left) and Regulon Stability Index RSI (right). (f) Integrated view: stable plateaux (blue, σ_1_ < 50th percentile) and phase boundaries (red contours, ⍰⍰C⍰ > 85th and 95th percentiles).

Regulon activity fields are constructed as weighted combinations of z-scored target gene fields. We defined five regulon programs relevant to skin biology: STAT3 (immune/inflammatory signaling), NF-κB (innate immunity), IRF7 (interferon response), KRT-diff (keratinocyte differentiation), and TRM (tissue-resident memory T cell signature). Target genes and weights were assigned based on established regulatory relationships (Methods). Of the 31 unique target genes specified across these five regulons, 12 were present in the Xenium panel (382 genes), with one to four targets available per regulon (Supplementary Table 1). Despite this incomplete coverage (39%), the coupling tensor — which integrates covariance across all regulon fields — proved robust to missing targets (see Limitations).

Applied to a Xenium human skin melanoma section (87,499 cells, 13.7 million transcripts), STRATA produced continuous regulon fields that recapitulated expected tissue biology (Fig. 1b). The KRT-diff field showed highest activity in the epidermal compartment, with the gradient vector field oriented perpendicular to the epidermal surface — consistent with the known basal-to-superficial differentiation axis of the epidermis (Fig. 1c). The Laplacian ⍰^2^φ(KRT-diff) was negative at the epidermal surface (convergent differentiation signal) and positive at the dermal–epidermal junction (DEJ; divergent signal), matching the known source-sink dynamics of keratinocyte differentiation. The immune regulon fields (STAT3, NF-κB, IRF7) showed enrichment in the peritumoral stroma, while TRM was concentrated at the tumor–stroma interface. The complete Layer 1 pipeline processed 13.7 million transcripts in 17.7 seconds on a single T4 GPU.

### The coupling tensor reveals local co-regulation geometry

The second layer of STRATA quantifies how regulon programs co-vary locally in space (Fig. 1d). At each tissue position s, we compute a coupling tensor C(s) — a P × P symmetric matrix (P = number of regulons) whose entries represent the local covariance between regulon fields within a Gaussian window of radius δ (default 100 μm). The coupling tensor captures the local co-regulation structure: which transcription factor programs rise and fall together at each tissue position.

From C(s) we derive two summary quantities. The coupling strength ⍰C⍰_F is the Frobenius norm of the tensor, measuring the total magnitude of local co-regulation. The effective dimensionality d_eff = exp(−Σ p_i log p_i), where p_i are the normalized eigenvalues of C(s), quantifies the complexity of the local coupling structure. A position where all regulon programs co-vary along a single axis has d_eff ≈ 1, while a position with independent regulatory variation in all directions has d_eff ≈ P.

Across the melanoma skin section, coupling strength was highest at tissue interfaces — the DEJ, the tumor margin, and perivascular niches — where multiple regulatory programs converge (Fig. 1d, left). The effective dimensionality was relatively uniform (mean d_eff = 3.01 across the tissue), indicating that approximately three independent modes of regulatory co-variation exist at most tissue positions with five regulon inputs. Regions of reduced d_eff (≈ 2) appeared within the tumor core, suggesting that the tumor microenvironment constrains regulatory variation to fewer independent modes.

### Phase boundaries mark where regulatory logic changes

We define coupling phase boundaries as positions where the coupling tensor changes rapidly in space: ⍰⍰C⍰ = [Σ_{i,j} (∂C_{ij}/∂x)^2^ + (∂C_{ij}/∂y)^2^]^{1/2}. These are not expression boundaries (where individual gene levels change) but co-regulation boundaries (where the *relationship* between regulatory programs changes). We hypothesized that phase boundaries should correspond to histologically defined tissue interfaces, because the co-regulation logic of keratinocytes differs fundamentally from that of fibroblasts or immune cells (Conjecture 5.1 in the STRATA mathematical framework).

To test this, we overlaid phase boundary contours onto H&E histology of the same tissue section (Fig. 2a,b). Phase boundaries at the 85th and 95th percentiles of ⍰⍰C⍰ aligned with the DEJ, the tumor–stroma interface, and boundaries between papillary and reticular dermis. This correspondence was quantified in two independent ways. First, the coupling gradient magnitude correlated with the KRT-diff field gradient (Pearson r = 0.3196, P < 10^−10^; Fig. 2d), confirming that positions of rapid coupling change coincide with the keratinocyte differentiation boundary. Second, the coupling gradient correlated more strongly with σ_1_ (r = 0.5094, P < 10^−10^), the maximum principal stretch from the Jacobian SVD (see below), providing internal mathematical consistency: two independent geometric quantities — one derived from the coupling tensor, the other from the regulon Jacobian — converge on the same spatial structures.

**Figure 2.**
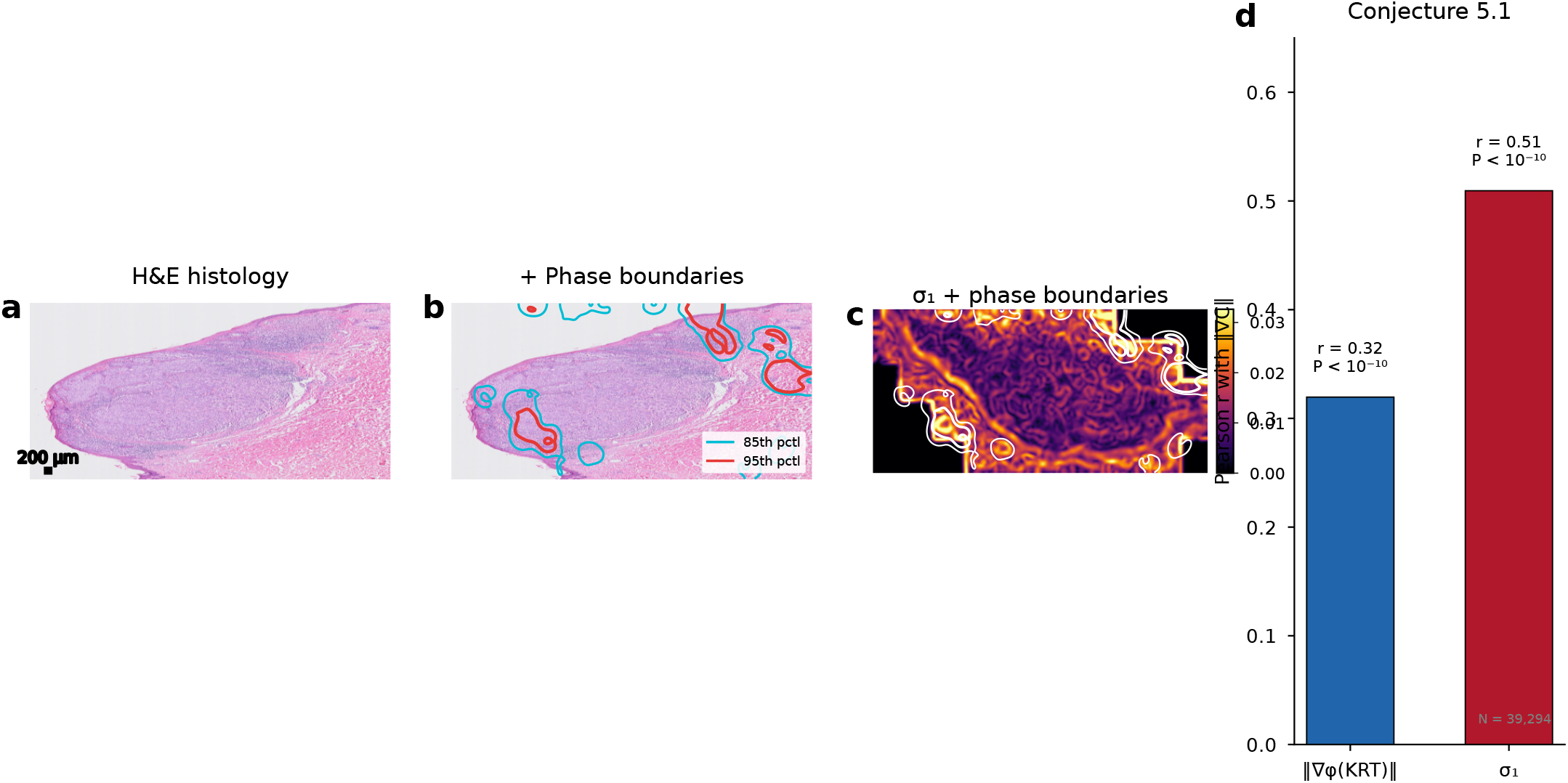
Phase boundaries correspond to histological tissue architecture (Conjecture 5.1). (a) H&E histology of the melanoma skin section. (b) H&E with overlaid phase boundary contours (cyan: 85th percentile; red: 95th percentile of ⍰⍰C⍰). Phase boundaries track the dermal–epidermal junction, tumor margins, and dermal compartment boundaries. (c) Maximum principal stretch σ_1_ heatmap with phase boundary contours (white). (d) Quantification: Pearson correlation between ⍰⍰C⍰ and ⍰⍰φ(KRT-diff) ⍰ (r = 0.32, P < 10^−10^) and between ⍰⍰C⍰ and σ_1_ (r = 0.51, P < 10^−10^). N = 39,294 spatial grid points.

### The Regulon Stability Index identifies rigid and labile regulatory configurations

The third layer of STRATA computes the spatial Jacobian J(s) of the regulon field map — the P × 2 matrix whose rows are the spatial gradients of each regulon field (Fig. 1e). The SVD of J(s) yields two principal stretches: σ_1_ (maximum) and σ_2_(minimum). The maximum principal stretch σ_1_ quantifies the rate at which the regulon configuration changes along the direction of greatest variation; it is large at tissue interfaces where regulatory programs transition sharply, and small within homogeneous tissue compartments. The Regulon Stability Index RSI = σ_2_/σ_1_ measures the isotropy of this change: RSI ≈ 1 indicates that the configuration changes equally in all directions (isotropic instability), while RSI ≈ 0 indicates change is confined to a single axis (anisotropic transition, characteristic of a sharp boundary).

Together, σ_1_ and RSI partition the tissue into stable plateaux (low σ_1_, high RSI — positions where the regulon configuration is locally constant) and phase boundaries (high σ_1_, low RSI — positions where it transitions sharply along one direction) (Fig. 1f). The stable plateaux correspond to histologically uniform tissue compartments; the phase boundaries correspond to tissue interfaces. This partitioning emerges purely from the mathematics of the regulon fields, without any reference to histology or cell type labels.

### Conjecture 5.1: quantitative validation against tissue architecture

We formally tested the correspondence between STRATA-derived phase boundaries and histological tissue architecture (Fig. 2). H&E histology was available for the same tissue section as the Xenium data, enabling direct spatial overlay. Phase boundary contours (95th percentile of ⍰⍰C⍰) traced the DEJ, tracked the tumor–stroma boundary, and delineated the transition between papillary and reticular dermis (Fig. 2b). The σ_1_ landscape showed high values precisely at these same interfaces (Fig. 2c).

Quantitatively, the spatial correlation between the coupling gradient ⍰⍰C⍰ and the KRT-diff gradient ⍰⍰φ(KRT-diff) ⍰ was r = 0.3196 (P < 10^−10^; N = 39,294 spatial grid points). While moderate in magnitude, this correlation is notable because it compares a multivariate geometric quantity (the coupling tensor gradient, integrating information across all five regulons) with a univariate field gradient (KRT-diff alone). The correlation with σ_1_ was substantially stronger (r = 0.5094, P < 10^−10^), indicating that the full Jacobian SVD — which captures the aggregate rate of change across all regulon fields — is a more sensitive marker of tissue architecture than any single regulon gradient.

These correlations were robust to parameter choice. Across 12 combinations of KDE bandwidth (20, 40, 60, 80 μm) and coupling window radius (50, 100, 150 μm), r(⍰⍰C⍰, ⍰⍰φ(KRT-diff)⍰) ranged from 0.168 to 0.371, and r(⍰⍰C⍰, σ_1_) ranged from 0.349 to 0.528, remaining significant in all cases (Supplementary Fig. 2). The default parameters (bandwidth 40 μm, δ = 100 μm) were chosen based on biologically motivated length scales (approximately two cell diameters and a local neighborhood radius, respectively), not optimized for these correlations.

### Melanoma homogenizes the coupling landscape

Having established that STRATA phase boundaries correspond to tissue architecture, we asked how melanoma affects the coupling landscape. Downloading a separate normal skin section for comparison would introduce batch effects and confound biological with technical variation. Instead, we exploited the natural tissue architecture of the section itself: the upper portion (y < 1458 μm) contains epidermis and superficial dermis with minimal tumor involvement, while the lower portion (y ≥ 1458 μm) contains the deep dermis and tumor mass. By running the full STRATA pipeline independently on each zone, we obtained an internally controlled comparison that isolates the effect of tumor presence on coupling architecture.

The results revealed an unexpected pattern (Fig. 3, Table 1). Mean coupling strength was nearly identical between zones (⍰C⍰_F: 0.4404 epidermal vs. 0.4045 tumor, ratio 0.92). Mean effective dimensionality was virtually unchanged (d_eff: 3.01 vs. 2.93). Mean σ_1_ was indistinguishable (0.0135 vs. 0.0136, ratio 1.01). By every measure of *mean* coupling, the tumor microenvironment was essentially normal.

**Figure 3.**
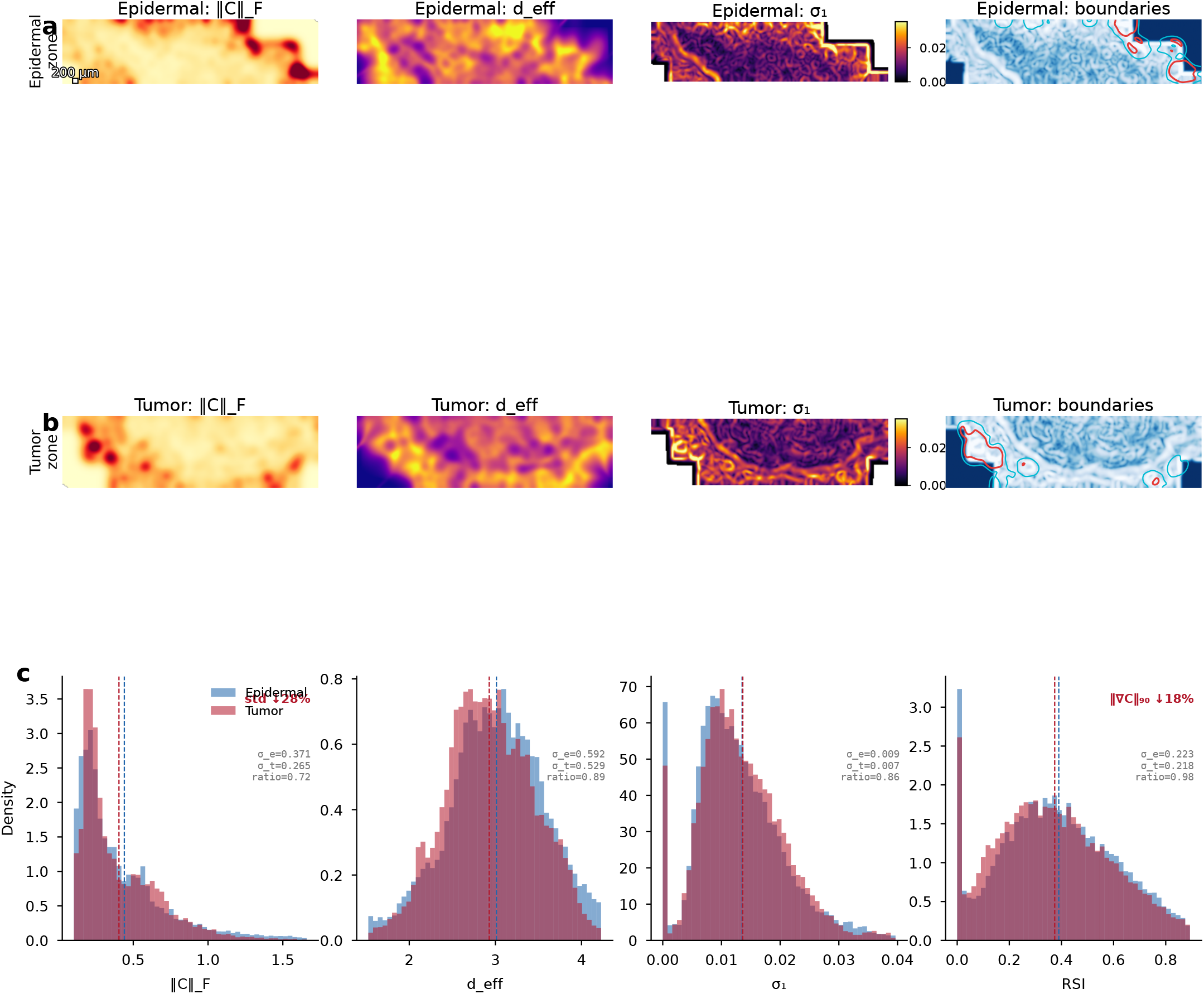
Melanoma homogenizes the coupling landscape. Top row: epidermal zone (y < 1458 μm) showing ⍰C⍰_F, d_eff, σ_1_, and plateaux/boundaries. Middle row: tumor zone (y ≥ 1458 μm), same metrics with shared color scales. Bottom panels: overlaid histograms comparing distributions between zones. Coupling strength, effective dimensionality, and principal stretch have nearly identical means between zones, but the tumor zone shows narrower distributions: coupling standard deviation decreases 28% and 90th percentile phase boundary intensity decreases 18%. Melanoma preserves coupling but erases its spatial organization.

**Table 1:**
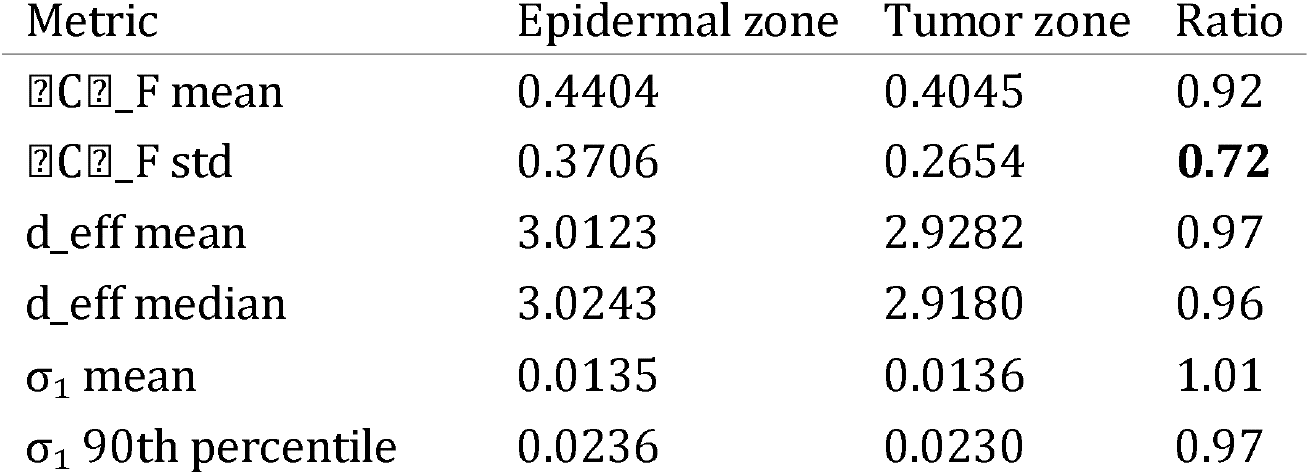
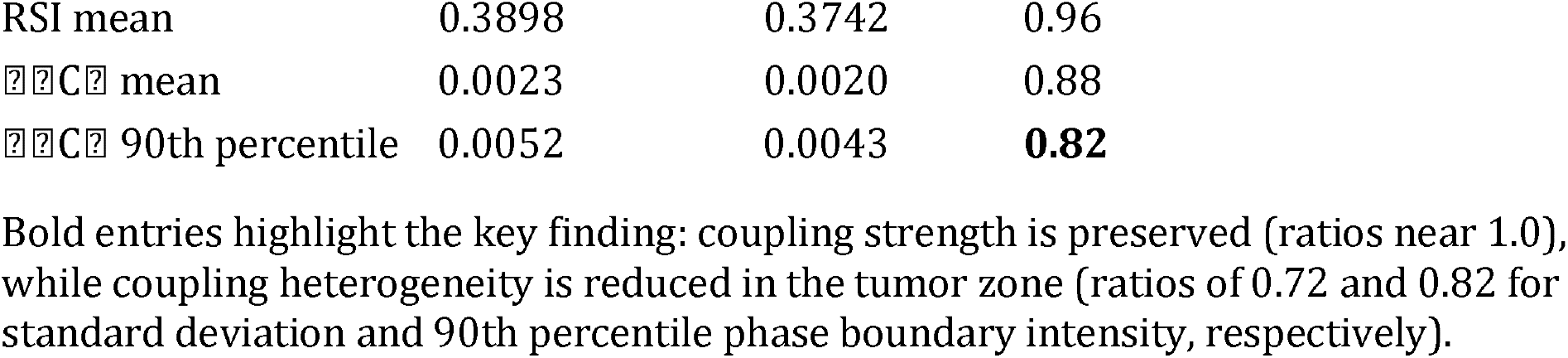
Coupling architecture comparison between epidermal and tumor zones.

The difference emerged in the *spatial heterogeneity* of coupling. The standard deviation of ⍰C⍰_F decreased 28% in the tumor zone (0.3706 to 0.2654). The 90th percentile of the phase boundary gradient ⍰⍰C⍰ decreased 18% (0.0052 to 0.0043). The tumor zone distributions were consistently narrower than the epidermal zone distributions across all metrics (Fig. 3, bottom panels). Melanoma does not destroy regulon coupling — it homogenizes the coupling landscape, erasing the structured spatial heterogeneity that characterizes organized tissue.

This finding is consistent with the known biology of melanoma invasion. As tumor cells infiltrate and replace normal tissue architecture, the structured layering of epidermis, DEJ, papillary dermis, and reticular dermis is progressively obliterated. In the STRATA framework, this architectural loss manifests not as reduced coupling strength but as reduced coupling variance — the tissue loses its organized heterogeneity while maintaining similar average regulatory co-variation. Standard clustering-based analysis would detect different cell type proportions between zones; STRATA reveals how the tumor reshapes the continuous regulatory geometry of tissue.

## Discussion

We have introduced STRATA, a differential-geometric framework that transforms spatial transcriptomics data from discrete cell catalogues into continuous regulon field analyses. Three core contributions emerge from this work.

First, the coupling tensor provides a fundamentally new quantity for spatial transcriptomics analysis. Existing methods characterize *what* is expressed at each tissue position (gene expression, cell types, pathway scores). The coupling tensor characterizes *how* regulatory programs relate to each other at each position — the local co-regulation geometry. This distinction matters because tissue function depends not on which programs are active in isolation, but on how they are coordinated. Two positions with identical mean expression of five regulons may have completely different coupling structures — one with all programs varying independently, another with tight co-regulation along a single axis. STRATA distinguishes these cases.

Second, phase boundaries represent a new class of spatial feature. Expression boundaries — where individual gene levels change — have been identified by various spatial methods. Phase boundaries are qualitatively different: they mark positions where the *relationship* between programs changes, even if individual expression levels do not. The DEJ is a phase boundary not primarily because gene expression changes there (although it does), but because the co-regulation logic of keratinocytes is fundamentally different from that of fibroblasts. STRATA captures this distinction.

Third, the melanoma coupling homogenization finding illustrates the kind of insight that emerges from field-theoretic analysis but is invisible to cell-centric approaches. The standard narrative would be that melanoma disrupts tissue — and indeed mean expression, cell type composition, and pathway scores all change in the tumor microenvironment. But STRATA reveals a more nuanced picture: the tumor preserves coupling strength while erasing coupling heterogeneity. The tissue is not less regulated; it is more uniformly regulated. This homogenization may reflect the tumor’s imposition of a dominant microenvironmental program that overrides the position-dependent regulatory diversity of normal skin architecture. Whether coupling homogenization is a general feature of solid tumors — and whether its degree correlates with invasive potential — are questions that STRATA is uniquely positioned to address across cancer types and spatial transcriptomics platforms.

Several limitations should be noted. The current implementation uses a fixed set of regulon definitions based on literature-curated target genes; data-driven regulon inference from the spatial data itself (for example, via spatial extensions of SCENIC or GRNBoost) could improve both coverage and specificity. The five regulons analyzed here were chosen for biological relevance to skin, and the framework’s behavior with larger regulon sets (tens to hundreds) remains to be characterized. The gene panel constraint (12 of 31 unique target genes available in the Xenium panel) limits the precision of individual regulon fields — particularly for IRF7, where only one target (ISG15) was available — although the coupling tensor, which integrates across regulons, appeared robust to this incompleteness. Finally, the current analysis is two-dimensional; extension to three-dimensional tissue reconstructions from serial sections is mathematically straightforward but computationally demanding.

STRATA is implemented as an open-source Python package (github.com/jengweitjiu/STRATA) with a complete reproducible notebook that processes 13.7 million transcripts in under 20 seconds on a standard GPU. The framework is platform-agnostic and applicable to any spatial transcriptomics data that provides transcript-level coordinates, including Xenium, MERFISH, CosMx, and Stereo-seq. By moving spatial transcriptomics analysis from cell cataloguing to continuous field theory, STRATA opens a new axis of investigation into the regulatory architecture of tissue in health and disease.

## Methods

### Data

We used the 10x Genomics Xenium Human Skin Gene Expression dataset (melanoma, FFPE), publicly available under CC BY 4.0 license. The dataset comprises 87,499 cells and 13,728,036 transcripts across 382 genes (282 pre-designed + 100 custom add-on panel) from a formalin-fixed paraffin-embedded human skin melanoma section. Transcript-level data (x, y coordinates and gene identity) were obtained as a Parquet file from the 10x Genomics website. Matched H&E histology was available as an OME-TIFF file.

### Regulon definitions

Five regulon programs were defined based on established transcriptional regulatory relationships in skin biology:

#### STAT3 regulon

Target genes (weights): SOCS3 (1.0), BCL2L1 (0.9), MYC (0.85), JUNB (0.7), VEGFA (0.7), MCL1 (0.85). Available in panel: MYC, VEGFA (2 of 6).

#### NF-κB regulon

Target genes (weights): NFKBIA (1.0), TNFAIP3 (0.95), CXCL8 (0.9), IL1B (0.9), CCL2 (0.85), ICAM1 (0.8). Available in panel: CCL2, ICAM1 (2 of 6).

#### IRF7 regulon

Target genes (weights): ISG15 (1.0), MX1 (0.95), IFIT1 (0.9), IFI44L (0.85), OAS1 (0.85), IFI6 (0.8), BST2 (0.75). Available in panel: ISG15 (1 of 7). Although only one target was available, ISG15 is a canonical type I interferon-stimulated gene with high dynamic range, and the resulting field recapitulated expected peritumoral enrichment.

#### KRT-diff regulon

Target genes (weights): KRT14 (−1.0), KRT5 (−0.9), KRT10 (1.0), KRT1 (0.9), FLG (0.8), LOR (0.7). Available in panel: KRT14, KRT10, KRT1 (3 of 6). Negative weights for basal keratins (KRT14, KRT5) encode the differentiation axis from basal (negative) to suprabasal (positive) keratinocyte identity.

#### TRM regulon

Target genes (weights): CD69 (1.0), ITGAE (1.0), ZNF683 (0.9), CD8A (0.7), CD8B (0.7), GZMB (0.6). Available in panel: CD69, ITGAE, CD8A, CD8B (4 of 6).

### Layer 1: Regulon field construction

For each gene g, we computed a transcript density field ρ_g(s) on a regular grid (resolution Δ = 20 μm) by binning transcript counts and smoothing with a Gaussian kernel of bandwidth h = 40 μm. Fields were log-normalized: x_g(s) = log(ρ_g(s)/Σ_g ρ_g(s) × 10^4^+ 1), then z-scored: z_g(s) = (x_g(s) − μ_g) / σ_g. Regulon fields were computed as: 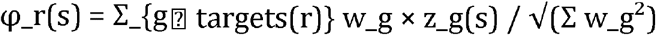, where w_g are the regulon weights. Spatial gradients ⍰φ and Laplacians ⍰^2^φ were computed using central finite differences on Gaussian-smoothed fields (σ_smooth = 1.5 grid cells).

### Layer 2: Coupling tensor and phase boundaries

The coupling tensor C(s) is a P × P symmetric matrix at each tissue position, defined as: C_{ij}(s) = G_δ * (φ_i − G_δ * φ_i)(φ_j − G_δ * φ_j), where G_δ denotes Gaussian smoothing with window radius δ = 100 μm and * denotes convolution. This computes the local covariance between regulon fields within a Gaussian neighborhood. The coupling strength is 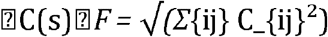. The effective dimensionality is d_eff(s) = exp(−Σ_k p_k log p_k), where p_k = λ_k / Σ λ_k are the normalized eigenvalues of C(s). Phase boundaries are defined as 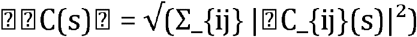, computed after Gaussian smoothing (σ = 2 grid cells) of each tensor component.

### Layer 3: Jacobian SVD and Regulon Stability Index

The regulon Jacobian J(s) is the P × 2 matrix whose rows are the spatial gradients of each regulon field: J_{r,:}(s) = ⍰φ_r(s). The SVD J = UΣV^T yields principal stretches σ_1_ ≥ σ_2_≥ 0. The maximum principal stretch σ_1_(s) quantifies the maximum rate of regulon configuration change at position s. The Regulon Stability Index is RSI(s) = σ_2_(s) / σ_1_(s) ⍰ [0, 1]. Stable plateaux are identified as regions with σ_1_ below the 50th percentile; phase boundaries as regions with σ_1_ above the 85th percentile.

### Within-tissue comparison

The tissue section was split at the y-axis midpoint (y = 1458 μm) into an upper zone (epidermis and superficial dermis; 6.2 million transcripts) and a lower zone (deep dermis and tumor mass; 7.5 million transcripts). The complete STRATA pipeline (Layers 1–3) was run independently on each zone with identical parameters. Coupling metrics (⍰C⍰_F, d_eff, σ_1_, RSI, ⍰⍰C⍰) were compared between zones using means, standard deviations, and percentile distributions.

### Parameter sensitivity analysis

Robustness was assessed across 12 parameter combinations: KDE bandwidth h ⍰ {20, 40, 60, 80} μm and coupling window δ ⍰ {50, 100, 150} μm. For each combination, the full pipeline was re-run and Conjecture 5.1 correlations (r(⍰⍰C⍰,⍰⍰φ(KRT-diff) ⍰) and r(⍰⍰C⍰, σ_1_)) were computed. Phase boundary spatial correlation with the reference parameter set (h = 40, δ = 100) was also quantified.

### H&E overlay

H&E histology was loaded from the OME-TIFF pyramid (level 4 for memory efficiency) and registered to the Xenium coordinate system using the manufacturer-provided transformation. Phase boundary contours were overlaid at the 85th and 95th percentiles of ⍰⍰C⍰.

### Software and reproducibility

All analyses were performed in Python using NumPy, SciPy, Matplotlib, and Pandas. The complete pipeline is provided as a Jupyter notebook (STRATA_Layer1_RealSkin_20260224_clean.ipynb) that runs in Google Colab on a T4 GPU with high RAM in under 20 seconds for the full dataset. Code is available at github.com/jengweitjiu/STRATA under an MIT license.

## Data availability

The Xenium Human Skin (Melanoma) dataset is available from 10x Genomics under CC BY 4.0 at https://www.10xgenomics.com/datasets/human-skin-preview-data-xenium-human-skin-gene-expression-panel-add-on-1-standard.

## Code availability

STRATA is available at https://github.com/jengweitjiu/STRATA under an MIT license. A complete reproducible notebook is included.

## Acknowledgements

None.

## Competing interests

The author declares no competing interests.

Bold entries highlight the key finding: coupling strength is preserved (ratios near 1.0), while coupling heterogeneity is reduced in the tumor zone (ratios of 0.72 and 0.82 for standard deviation and 90th percentile phase boundary intensity, respectively).

## Supplementary Figure legends

**Supplementary Figure 1. Synthetic tissue validation**. STRATA pipeline applied to a synthetic tissue with known ground truth regulon fields and planted interfaces, demonstrating recovery of phase boundaries at correct positions.

**Supplementary Figure 2. Parameter sensitivity analysis**. Heatmaps showing Conjecture 5.1 correlations across 12 parameter combinations (KDE bandwidth × coupling window). r(⍰⍰C⍰, ⍰⍰φ(KRT-diff) ⍰) ranges from 0.168 to 0.371; r(⍰⍰C⍰, σ_1_) ranges from 0.349 to 0.528. All correlations are significant (P < 10^−10^). Cross-parameter phase boundary correlation with reference set ranges from 0.341 to 0.933.

**Supplementary Figure 3. Gene availability analysis**. Assessment of target gene coverage: 12 of 31 unique target genes across five regulons are available in the Xenium panel (39% coverage), with one to four targets per regulon.

**Supplementary Table 1. Regulon target gene availability in the Xenium panel**. Complete listing of target genes, weights, and panel availability for all five regulons.

