## Supplementary figures and images for "STRATA: Spatial Regulon Field Theory Reveals Coupling Architecture of Human Skin and Its Homogenization in Melanoma"

### Supplemental Figure 1

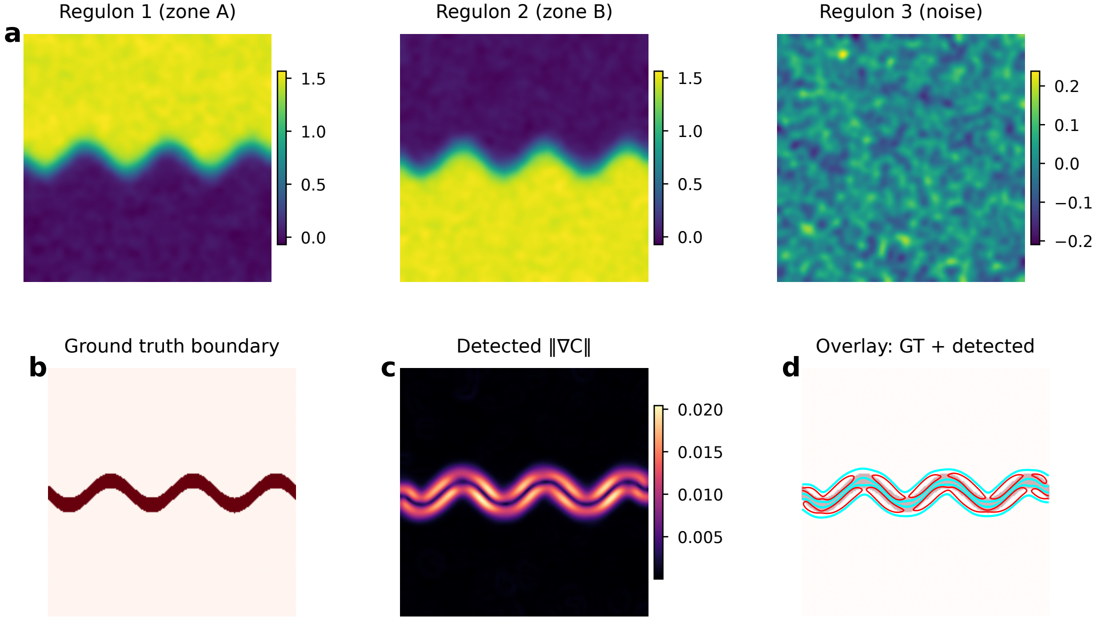

### Supplemental Figure 2

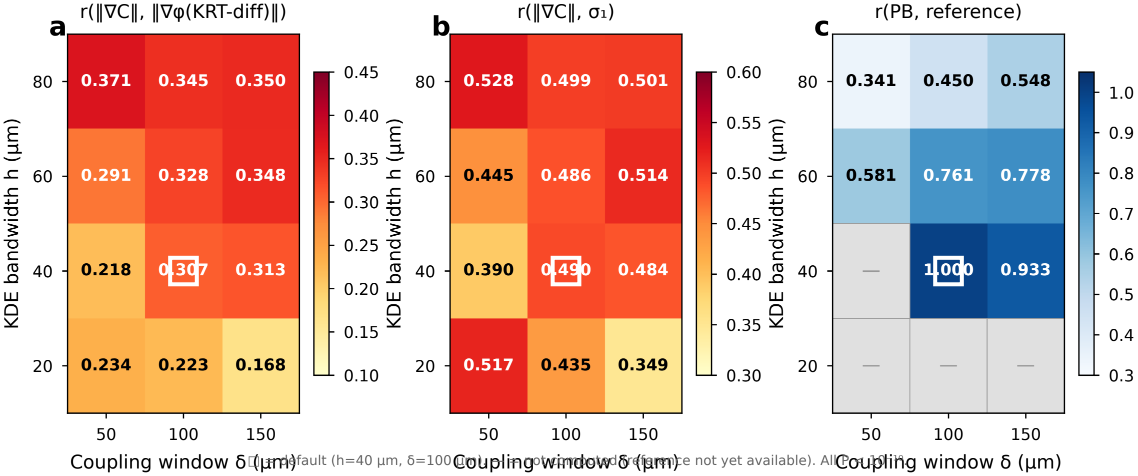

### Supplemental Figure 3

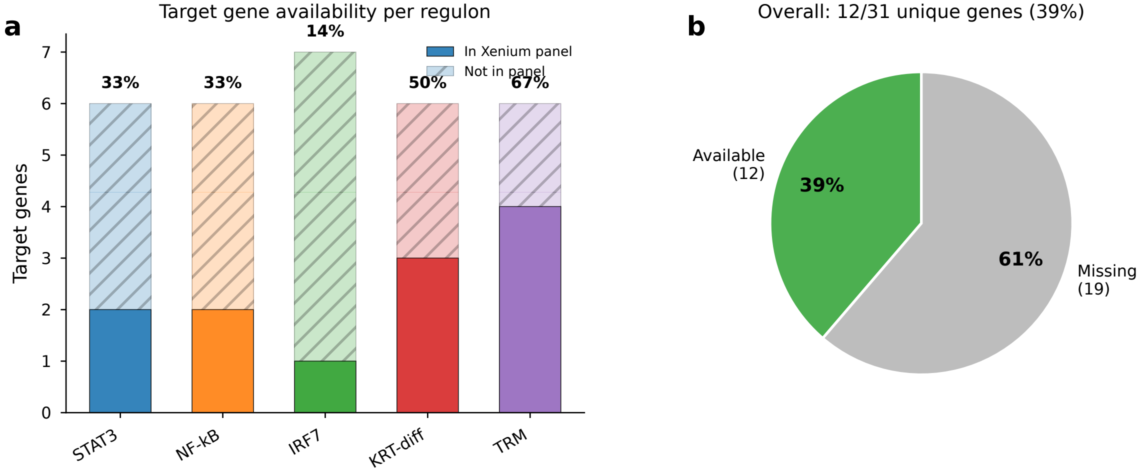
